# Stand structure as the proximate driver of endemic biodiversity and ecosystem functions in tropical mosaic landscapes

**DOI:** 10.1101/2024.03.20.582692

**Authors:** Rindrasoa Rajaonarimalala, Yevgeniya Korol, Aristide Andrianarimisa, Saskia Dröge, Thio Rosin Fulgence, Ingo Grass, Holger Kreft, Kristina Osen, Anjaharinony A. N. A. Rakotomalala, Bakolimalala Rakouth, Hery Lisy Tiana Ranarijaona, Romual Randriamanantena, Fanomezana Mihaja Ratsoavina, Lala Harivelo Raveloson Ravaomanarivo, Estelle Raveloaritiana, Dominik Schwab, Marie Rolande Soazafy, Teja Tscharntke, Annemarie Wurz, Dirk Hölscher, Dominic Andreas Martin

## Abstract

Transformation of forests into agricultural lands threatens biodiversity and ecosystem functions globally. In the biodiversity hotspot Madagascar, key ecosystem functions along with highly endemic flora and fauna are under threat. Comprehensive studies identifying proximate drivers are lacking, with no studies accounting for endemic species richness across multiple taxa. We assess how plot-scale stand structural attributes (basal area, leaf area index, diameter diversity, and tree species richness) and landscape-scale forest cover affect biodiversity (species richness of butterflies, ants, birds, amphibians, reptiles, herbaceous plants, and multidiversity) and ecosystem functions (soil organic carbon, predation rate, acoustic diversity index, and multifunctionality) in the tropical mosaic landscape of northeastern Madagascar. Across a prevalent land-use gradient in the region, we sampled five woody land-use types: old-growth forest, forest fragment, woody fallow, forest-derived and fallow-derived vanilla agroforests. Complexly structured stands with a larger basal area and higher tree species richness promoted greater endemic multidiversity and certain ecosystem functions. Landscape-scale forest cover significantly favored, particularly, endemic bird species richness. Our models explained 70% of the variance in endemic multidiversity, 26% in multidiversity, and 43% in multifunctionality. Significant effects directions of our predictors differed between taxa but were univocally positive for endemic species richness. Our findings highlight that a simplified stand structure, resulting from land-use change, substantially reduces endemic biodiversity and ecosystem functions in this historically forested region. Conservation actions should aim at maintaining complex stand structure in forests and agroforests, while restoration interventions should focus on re-building such structures.

## 1. Introduction

Tropical forests host a significant part of the world’s biodiversity (Barlow et al., 2018). Their preservation is challenged by widespread and ongoing deforestation and forest degradation (Newbold et al., 2015; Pendrill et al., 2022). Large-scale forest conversion for commodity production dominates in parts of South America and Southeast Asia, while subsistence-driven shifting cultivation is the main driver of forest loss in most of Africa, including Madagascar (Curtis et al., 2018). Malagasy forests have lost about 44% of their area since 1953 (Vieilledent et al., 2018), mostly due to shifting agriculture (Zaehringer et al., 2015). Associated biodiversity decline is alarming (Ralimanana et al., 2022) given the outstanding rates of endemism in the Malagasy flora and fauna (Antonelli et al., 2022) and the contribution of biodiversity to ecosystem functioning (Le Provost et al., 2022; Yan et al., 2023) and livelihoods (Blicharska et al., 2019). The preservation of ecosystem functions is crucial to ensure the multitude of benefits they provide, including provisioning, regulating, supporting, and cultural services (Garland et al., 2021).

Conventionally, conservation and restoration approaches in the humid tropics aim to conserve forests and high tree cover within mosaic landscapes (Arroyo-Rodríguez et al., 2020; Wright, 2010). While approaches may vary, they often focus on preserving or restoring trees, thereby affecting stand structure. Habitat complexity, describing within-habitat variability, often refers to the vegetation component of environmental heterogeneity (Stein and Kreft, 2014). Tree-based ecosystems derive their habitat complexity from the enhanced structural characteristics of trees, including their size variation, species diversity, and spatial distribution (Pommerening, 2002), hereafter called stand structure. Previous studies have revealed how relationships between stand structure and biodiversity (Atikah et al., 2021; Gebremichael et al., 2022; Penone et al., 2019; Storch et al., 2023), and between stand structure and ecosystem functions (Ali, 2019; Felipe-Lucia et al., 2018; Firn et al., 2007) are highly variable and context-specific. To achieve general conclusions, it is therefore crucial to study multiple taxa and functions. Here, multidiversity and multifunctionality, defined as generalized indices over several taxa and functions (Allan et al., 2014; Manning et al., 2018), can provide insight into biodiversity and ecosystem function responses (Ampoorter et al., 2020; Ouyang et al., 2023; Penone et al., 2019).

Conservation and restoration actions - if implemented on a larger scale - also affect landscape-scale forest cover. Forest-dependent species often require habitats at the landscape scale (Arroyo-Rodríguez et al., 2020) and landscape-scale forest cover is linked to higher seed dispersal by animals (San-José et al., 2020), pest regulation (Aristizábal and Metzger, 2019), and other ecosystem functions (Tolessa et al., 2017). Knowledge of the relationship between stand structure or forest cover and biodiversity and ecosystem functions may help to identify areas of potentially high endemic diversity using remote sensing data (Kacic and Kuenzer, 2022), for example, guiding conservation efforts towards priority areas in tropical landscapes under transition, such as in northeastern Madagascar. In addition, landscape rehabilitation or ecosystem restoration approaches by planting trees on degraded and formerly forested land may be better targeted towards structural features and increasing forest land cover in the landscape (Arroyo-Rodríguez et al., 2020).

Tropical landscapes are often rich in forest-dependent endemic species (Goodman and Benstead, 2005). Madagascar’s high level of endemism is due to its isolation and adaptive radiation (Antonelli et al., 2022; Goodman and Benstead, 2005; Vences et al., 2009), which means that small populations have evolved and adapted to specific environmental conditions. Indeed, the dynamic escarpments along the eastern coast of Madagascar created barriers contributing to population isolation and speciation (Liu et al., 2024). Endemics are therefore vulnerable to habitat change (Bellard et al., 2014). A strong relationship between endemism and stand structure is expected, but a comprehensive knowledge of this relationship and diverse parameters is lacking. To our knowledge, no studies accounting for endemic species richness across multiple taxa exist so far. In addition, understanding how each taxon responds to stand structure and forest cover is also important for species conservation, as it enables us to determine which taxa would benefit most.

In this study, we assess the connection between multiple plot-scale stand structural attributes and landscape-scale forest cover with biodiversity, as well as ecosystem functions, in the mosaic landscape of northeastern Madagascar. The stand structural attributes include basal area, leaf area index (LAI), diameter diversity, and tree species richness. Based on findings by McElhinny et al. (2005), we included tree species richness as a stand structural measure. In particular, we focus on the plot-scale stand structure of tree-based land-use types across a forest degradation gradient: old-growth forest, forest fragment, and forest-derived vanilla agroforest, and a restoration gradient: woody fallow and fallow-derived vanilla agroforest (Martin et al., 2020; Fig. 1). We test the following hypotheses: (a) Species richness of six taxonomic groups (butterflies, ants, birds, amphibians, reptiles, and herbaceous plants), as well as multidiversity, consistently increases with complex stand structure and greater landscape-scale forest cover. We expect a stronger effect of stand structure on endemic species richness; (b) Ecosystem functions (soil organic carbon (SOC) stock, predation rate, and acoustic diversity index), as well as multifunctionality, are promoted by complex stand structure and higher landscape-scale forest cover.

**Fig. 1.**
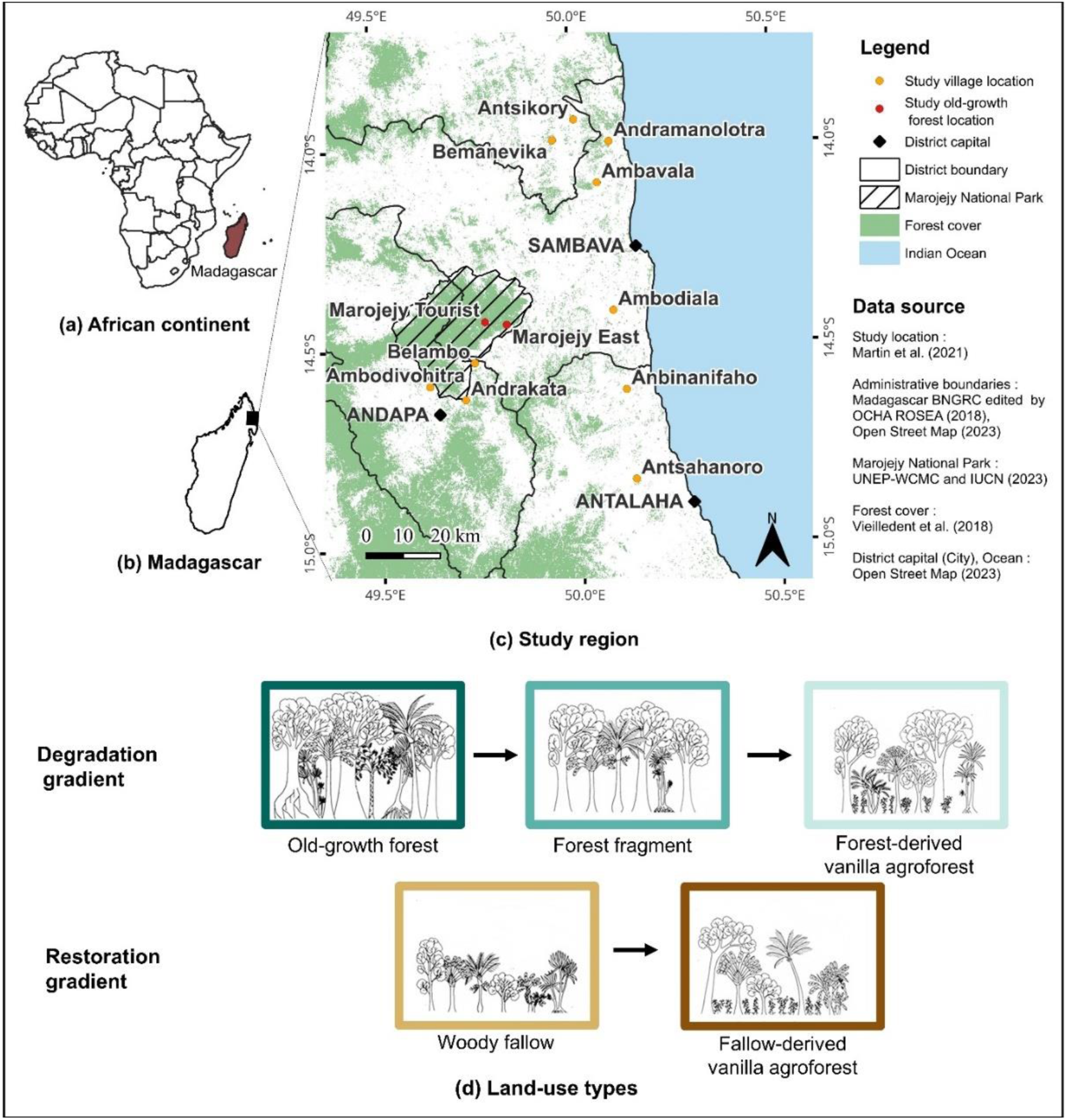
Study region in northeastern Madagascar. a) Continent of Africa and the island of Madagascar in East Africa. (b) Island of Madagascar. (c) Overview of the study region. Data collection (stand structure, biodiversity, and ecosystem function) in two sites of Marojejy National Park (red dots, with 10 replicates for old-growth forest) and in the surroundings of 10 villages (yellow dots, with 10 replicates for forest fragment, woody fallow, and forest-derived vanilla agroforest, and 18 replicates for fallow-derived vanilla agroforest; refer to Martin et al. (2022) and Wurz et al. (2022b) for more details). (d) Land-use gradients in the region (Martin et al., 2020). Artworks from Soazafy et al. (2021).

## 2. Material and methods

### 2.1. Study region

The study region is situated in northeastern Madagascar (Fig. 1). The climate is tropical, with a mean annual temperature of 23.9°C and an annual rainfall of 2 258 mm in the study region (58 plots; Karger et al., 2017; Osen et al., 2021). Moist forest is the natural vegetation type in this region (Vieilledent et al., 2016) and nowadays mainly persists within protected areas. The region is extremely rich in biodiversity, with high endemism (Martin et al., 2022; Wurz et al., 2022b).

### 2.2. Inventory design

From October 2017 to February 2019 (Martin et al., 2022), we sampled a total of 58 circular plots of 25 m radius (Fig. A1; 1 963.5 m^2^) in five prevalent woody land-use types in the region. We sampled old-growth forest at two sites in Marojejy National Park, the only large and continuous forest block (Fig. 1), forest fragment defined as regularly used but unburnt natural woodland embedded in the agricultural landscape, and woody fallow - fallow land where woody vegetation is regenerating after shifting hill rice cultivation. Moreover, we considered forest-derived vanilla agroforest, which is established in a forest by thinning understory vegetation, and fallow-derived vanilla agroforest - where vanilla is planted on fallow land (Osen et al., 2021; Wurz et al., 2022a).

### 2.3. Plot-scale stand structure

On every plot, we measured diameter at breast height (DBH) and determined all species of trees with DBH ≥8 cm (Fig. A1), including arborescent herbs and palms, tree ferns, but excluding lianas (Osen et al., 2021; Soazafy et al., 2021). From these measured variables, we derived basal area (m^2^/ha), diameter diversity (effective number of DBH classes; Table A1; Ehbrecht et al., 2019), and tree species richness (number of tree species per plot).

We assessed LAI (m^2^_leaf area_/m^2^_ground_) using a LI-COR LAI-2200C Plant Canopy Analyzer (LI-COR, Lincoln, USA), with two sensors using 90° angle view caps and setting up at a height of 2.4 m using a tripod. One sensor was inside the plot, collecting data below the canopy, while the other sensor in the surrounding gaps gathered data above the canopy. We computed the average value per plot from 100 readings (Osen et al., 2021).

### 2.4. Landscape-scale forest cover

We used a binary forest cover layer with a 30 m resolution from the year 2017 (Vieilledent et al., 2018). We considered the percentage of forest cover within a 250 m radius around each plot center (Fig. A1), representing a compromise for mobile and immobile taxa and avoiding radius overlap since the minimum distance between plots was 250 m (Martin et al., 2021a).

### 2.5. Biodiversity

We considered six different taxa: butterflies, ants, birds, amphibians, reptiles, and herbaceous plants. The sampling positions within plots are available in Fig. A1. To quantify biodiversity, we used total species richness per plot and endemic species richness per plot.

For butterflies, we set up eight traps with fermented bananas for 24 h in the interior of the plot. Additionally, we captured butterflies with time-standardized netting by walking 30 min slowly and steadily. We dried butterflies and afterwards determined the species in the laboratory. We did not include moths (Wurz et al., 2022a).

For ants, we sampled them in five stations. In each station, we installed a pitfall trap consisting of a plastic cup filled at one-third with 70% ethanol and a few drops of unscented soapy liquid for two days, and two bait traps 5 cm apart containing sugar and sardine for 30 min. We stored ant specimens in plastic vials filled with 70% ethanol for identification in the laboratory (Rakotomalala et al., 2021).

For birds, we carried out two 40-min point counts per plot. We started point counts no later than 8:15 a.m. with two observers and when weather conditions were favorable (no wind; no rain). We excluded birds outside the 25 m radius plot or overflying from the analysis (Martin et al., 2021a).

For amphibians and reptiles, we conducted time-standardized surveys of herpetofauna species within each plot, through a 45-min walk by two persons during both day and night, with three replicates each. Two repetitions were conducted during the dry season and one during the wet season. We based species identification on morphological features or molecular identification using DNA, if necessary (Fulgence et al., 2022).

For herbaceous plants, we installed eight subplots of 2 m × 2 m in each plot to assess all vascular plants without apparent woody stems. We determined their scientific species names based on their morphological characteristics. In cases where identification was challenging in the field, we collected herbarium samples for further identification (Raveloaritiana et al., 2021).

We calculated plot-level multidiversity (Allan et al., 2014) as the mean of the normalized species richness of each taxon, using formulas (1) and (2), where *M_i_* is the multidiversity at each plot *i*, *y_ij_* is the normalized species richness of taxon *j* at plot *i, x_ij_* is the species richness of taxon *j* at plot *i*, and N is the total number of studied taxa.

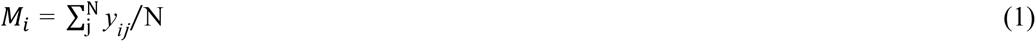

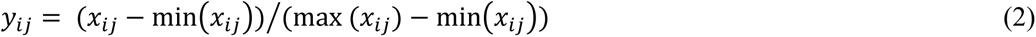

### 2.6. Ecosystem functions

Soil organic carbon stock indicates longer-term process rates (Manning et al., 2018) and soil is a substantial carbon reservoir on the planet (Lal, 2004). We extracted two mixed soil samples from the soil surface to 15 cm of depth with a corer of 5 cm diameter at eight different positions from each plot: the first from four cardinal directions and the second from intercardinal directions (Fig. A1). Afterward, we used a C/N elemental analyzer (Vario EL III, Elementar, Hanau, Germany) to measure the SOC value (mmol/g dry soil; Martin et al., 2022).

Predation rate is categorized as a regulating function, reflecting trophic interactions between organisms as natural biocontrol in ecosystems (Garland et al., 2021). For its estimation, we placed dummy plasticine caterpillars in all four cardinal axes (9 caterpillars arranged in a 3 m × 3 m grid in each axe, 36 caterpillars per plot, at one meter of height during 48 h, Fig. A1). We estimated the predation rate (%) by the percentage of the number of caterpillars attacked by vertebrate and invertebrate predators versus the number of retrieved dummy caterpillars (Schwab et al., 2021).

Soundscapes hold ecological and cultural value. They convey natural processes, habitat richness and quality (Pavan, 2017; Schafer, 1977), and contribute to human well-being (Pijanowski et al., 2011). For acoustic diversity, we deployed self-built solo audio recorders (Whytock and Christie, 2017) equipped with two microphones to register soundscapes inside the plot (Fig. A1). We used the R package *soundecology* (Villanueva-Rivera and Pijanowski, 2018) to determine the acoustic diversity index (ADI; Dröge et al., 2021).

We calculated multifunctionality in the same way as multidiversity, i.e., as the average of the normalized value of each ecosystem function. We applied the formulas (1) and (2), where *M_i_* is the multifunctionality at each plot *i*, *y_ij_* is the normalized value of ecosystem function *j* at plot *i*, *x_ij_* is the value of ecosystem function *j* at plot *i*, and N is the total number of studied ecosystem functions.

### 2.7. Statistical analysis

We performed all statistical analyses using R version 4.3.2 (R Core Team, 2023). Our study focused on determining the relationship of plot-scale stand structure as well as landscape-scale forest cover with biodiversity (species richness of six taxa and multidiversity) and ecosystem functions (three functions and multifunctionality). We considered five predictors: basal area, LAI, diameter diversity (effective number of DBH), tree species richness, and landscape-scale forest cover as predictors following preliminary analyses. The latter consisted of checking collinearity between predictors (Fig. A2 and Table A2) and exploring correlations between all potential predictors and response variables (Table A3). We assessed the strength, direction, and form of the relationship using a scatterplot and determining Spearman correlation coefficient r because of the nonlinear relationship between predictors and response variables and the non-normality of their distribution.

To examine the collective effect of all predictors, we fitted Generalized Linear Models implemented in the R package *glmmTMB* (Brooks et al., 2017). We used a Gaussian distribution for continuous response variables, a Poisson distribution for count data, and a negative binomial distribution in the case of overdispersion. We utilized the R packages *DHARMa* (Hartig, 2022) and *performance* (Lüdecke et al., 2021) to assess collinearity, residuals (uniformity, dispersion, and heterogeneity), singularity and convergence of the models, and zero inflation with count data as a response (Table A2). We extracted the coefficient of determination R² of each model using the r.squaredGLMM function in the R package *MuMIn* (Bartoń, 2023). We conducted hierarchical partitioning of R² to evaluate the variance in biodiversity and ecosystem functions explained by our predictors individually using the R package *glmm.hp* (Lai et al., 2022). We plotted all graphics using the R package *ggplot2* (Wickham, 2016).

## 3. Results

### 3.1. Determinants of species richness

Correlation analyses showed generally significant positive relationships between predictors and species richness of three of the six taxa (birds, reptiles, amphibians), and multidiversity (Fig. 2 and Fig. A3). Among the investigated plot-scale stand structural attributes, basal area and tree species richness showed the strongest positive correlation with reptile species richness (r= 0.55, p < 0.001; r= 0.55, p < 0.001), and LAI with amphibian species richness (r = 0.46, p < 0.001; Fig. A3). LAI and butterfly species richness were negatively correlated (r= −0.51, p < 0.001; Fig. A3). Multidiversity had weak correlations with stand structural attributes (Fig. 2). Landscape-scale forest cover had the strongest positive effect on bird species richness (r = 0.56, p < 0.001; Fig. A3) and multidiversity (r = 0.54, p < 0.001; Fig. 2).

**Fig. 2.**
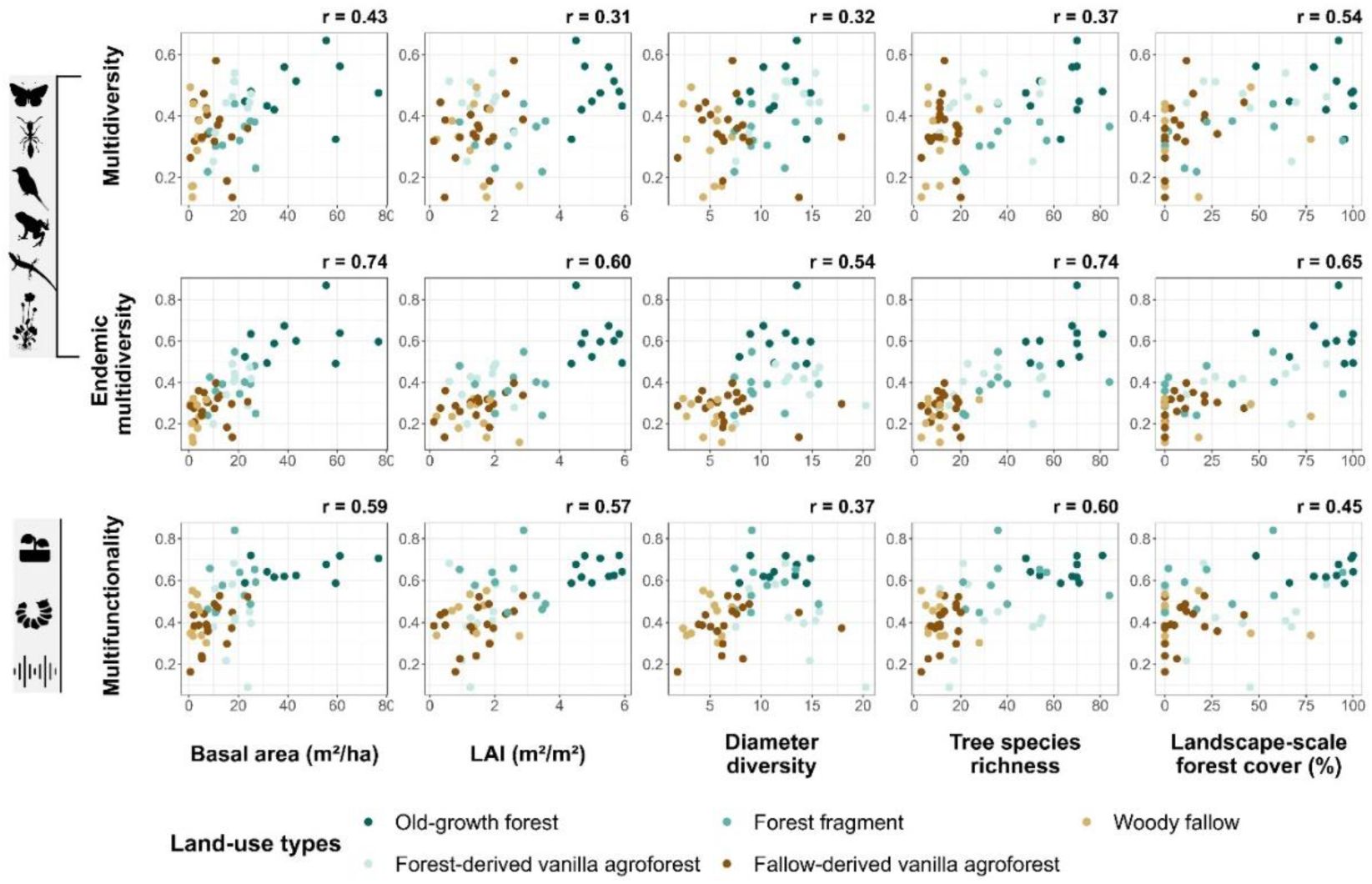
Correlation analyses between predictors and (endemic) multidiversity, as well as multifunctionality. Scatterplots are colored by land-use type. r is the Spearman correlation coefficient and significant correlation is in bold font (p < 0.05). Icons from phylopic.org and icons8.de (see Table A7 for attribution).

Full models considering all predictors in concert explained 26% of the variance in multidiversity (Table 1). The direction of the predictor-response relationship varied among taxa, half were generally positive and half negative (Table 2). On one hand, two predictors had a significant positive association with our response variables: tree species richness with reptile diversity (Est. = 0.23, SE = 0.07, p = 0.002), as well as landscape-scale forest cover with bird species richness (Est. = 0.17, SE = 0.07, p = 0.013; Table 2 and Table A4). On the other hand, two predictors were negatively associated with them: LAI with butterfly species richness (Est. = −0.26, SE = 0.09, p = 0.004), and also tree species richness with herbaceous plant richness (Est. = −0.20, SE = 0.09, p = 0.032; Table 2 and Table A4). None of the predictors were significant in determining multidiversity (Table 1).

**Table 1.** Effect of plot-scale stand structure and landscape-scale forest cover on biodiversity, endemic biodiversity, and ecosystem functions: GLM results. Significant predictors (p < 0.05) in bold font. Est.: estimate, SE: standard error, R²: coefficient of determination. More details about the model results and assessments are in Tables A2, A5, and A6.

| Predictors | | Intercept | Basal area | LAI | Diameter diversity | Tree species richness | Landscape-scale forest cover | $R^2$ |
| --- | --- | --- | --- | --- | --- | --- | --- | --- |
| Responses |  |  |  |  |  |  |  |  |
| **Multidiversity** | Est. | 0.38 | 0.02 | <0.01 | 0.01 | <0.01 | 0.03 | 0.26 |
|  | SE | 0.01 | 0.03 | 0.02 | 0.02 | 0.02 | 0.02 |  |
|  | z value | 30.19 | 0.76 | 0.09 | 0.35 | 0.20 | 1.69 |  |
| **Endemic Multidiversity** | Est. | 0.36 | **0.05** | 0.03 | <-0.01 | **0.04** | 0.02 | 0.70 |
|  | SE | 0.01 | **0.02** | 0.02 | 0.01 | **0.02** | 0.02 |  |
|  | z value | 34.08 | **2.31** | 1.67 | -0.23 | **2.41** | 1.29 |  |
| **Multi-functionality** | Est. | 0.48 | 0.05 | 0.04 | -0.03 | 0.05 | -0.01 | 0.43 |
|  | SE | 0.01 | 0.03 | 0.02 | 0.02 | 0.03 | 0.02 |  |
|  | z value | 32.23 | 1.58 | 1.52 | -1.38 | 1.76 | -0.26 |  |

**Table 2.** Direction of the relationship between predictors and response variables. Positive (+) and negative (−) indicate the direction of the relationship. The number x indicates the number of responses in a positive or negative relationship and the number between brackets represents the number of significant associations (p < 0.05). Details of model results and assessments are in Tables A2, A4, A5, and A6.

| Predictors | Basal area |  | LAI |  | Diameter diversity |  | Tree species richness |  | Landscape-scale forest cover |  |
| --- | --- | --- | --- | --- | --- | --- | --- | --- | --- | --- |
| Responses | + | - | + | - | + | - | + | - | + | - |
| **Species richness of six taxa for total species** | 3 (0) | 3 (0) | 4 (0) | 2 (1) | 2 (0) | 4 (0) | 3 (1) | 3 (1) | 4 (1) | 2 (0) |
| **Species richness of six taxa for endemic species** | 3 (2) | 3 (0) | 5 (0) | 1 (0) | 3 (1) | 3 (0) | 5 (1) | 1 (0) | 4 (1) | 2 (0) |
| **Three ecosystem functions** | 2 (1) | 1 (0) | 3 (0) | 0 (0) | 1 (0) | 2 (1) | 2 (1) | 1 (0) | 2 (0) | 1 (0) |

### 3.2. Determinants of endemic richness

Correlation analyses indicated positive associations between predictors and species richness of five to six taxa (except butterflies) and endemic multidiversity (Fig. 2 and Fig. A4). In terms of plot-scale stand structure, the strongest positive correlation occurred between basal area as well as tree species richness and endemic multidiversity (r = 0.74, p < 0.001; r = 0.74, p < 0.001; Fig. 2). Basal area and tree species richness also displayed a significant positive correlation with ant species richness (r = 0.72, p < 0.001; r = 0.73, p < 0.001; Fig. A4). Landscape-scale forest cover had the strongest positive correlation with endemic ant species richness (r = 0.67, p < 0.001) and endemic bird species richness (r = 0.66, p < 0.001; Fig. A4).

Full models including all predictors explained 70% of the variance in endemic multidiversity (Table 1). Our results showed only positive significant predictors in explaining endemic species richness (Table 2): basal area with endemic butterflies (Est. = 0.22, SE = 0.10, p = 0.030) and endemic birds (Est. = 0.32, SE = 0.12, p = 0.007), tree species richness with endemic reptiles (Est. = 0.24, SE = 0.08, P = 0.004), diameter diversity with endemic ants (Est. = 0.25, SE = 0.07, p < 0.001), and landscape-scale forest cover with endemic birds (Est. = 0.45, SE = 0.13, p < 0.001; Table A4). Basal area (Est. = 0.05, SE = 0.02, p = 0.021) and tree species richness (Est. = 0.04, SE = 0.02, p = 0.016) emerged as significant predictors of endemic multidiversity (Table 1).

### 3.3. Determinants of ecosystem functions

Correlation analyses revealed positive relationships between predictors and all studied ecosystem functions, including multifunctionality (Fig. 2 and Fig. A5). The strongest correlations were between tree species richness and acoustic diversity (r = 0.62, p < 0.001). Multifunctionality was positively correlated with tree species richness (r = 0.60, p < 0.001), basal area (r = 0.59, p < 0.001), and LAI (r = 0.57, p < 0.001; Fig. 2).

In fitted full models, predictors explained 43% of the variance in multifunctionality (Table 1). The direction of the predictor-response relationship differed across all three ecosystem functions (Table 2). Two predictors were positively associated with our response variables: basal area with SOC (Est. = 1.00, SE = 0.28, p < 0.001) and tree species richness with acoustic diversity index (Est. = 0.16, SE = 0.08, p = 0.049), whereas diameter diversity was negatively associated with SOC (Est. = −0.49, SE = 0.19, p = 0.012; Table A4).

### 3.4. Relative importance of plot-scale stand structure and landscape-scale forest cover

The relative importance of each predictor differed across taxa and individual ecosystem function. The total explanatory power of all predictors combined was particularly high for endemic taxa. Generally, basal area, LAI, and tree species richness explained a large proportion of the variance in endemic multidiversity and multifunctionality. Landscape-scale forest cover was important, especially for bird species richness, endemic bird species richness and endemic multidiversity. Diameter diversity had the smallest importance for all taxa except ants (Fig. 3).

**Fig. 3.**
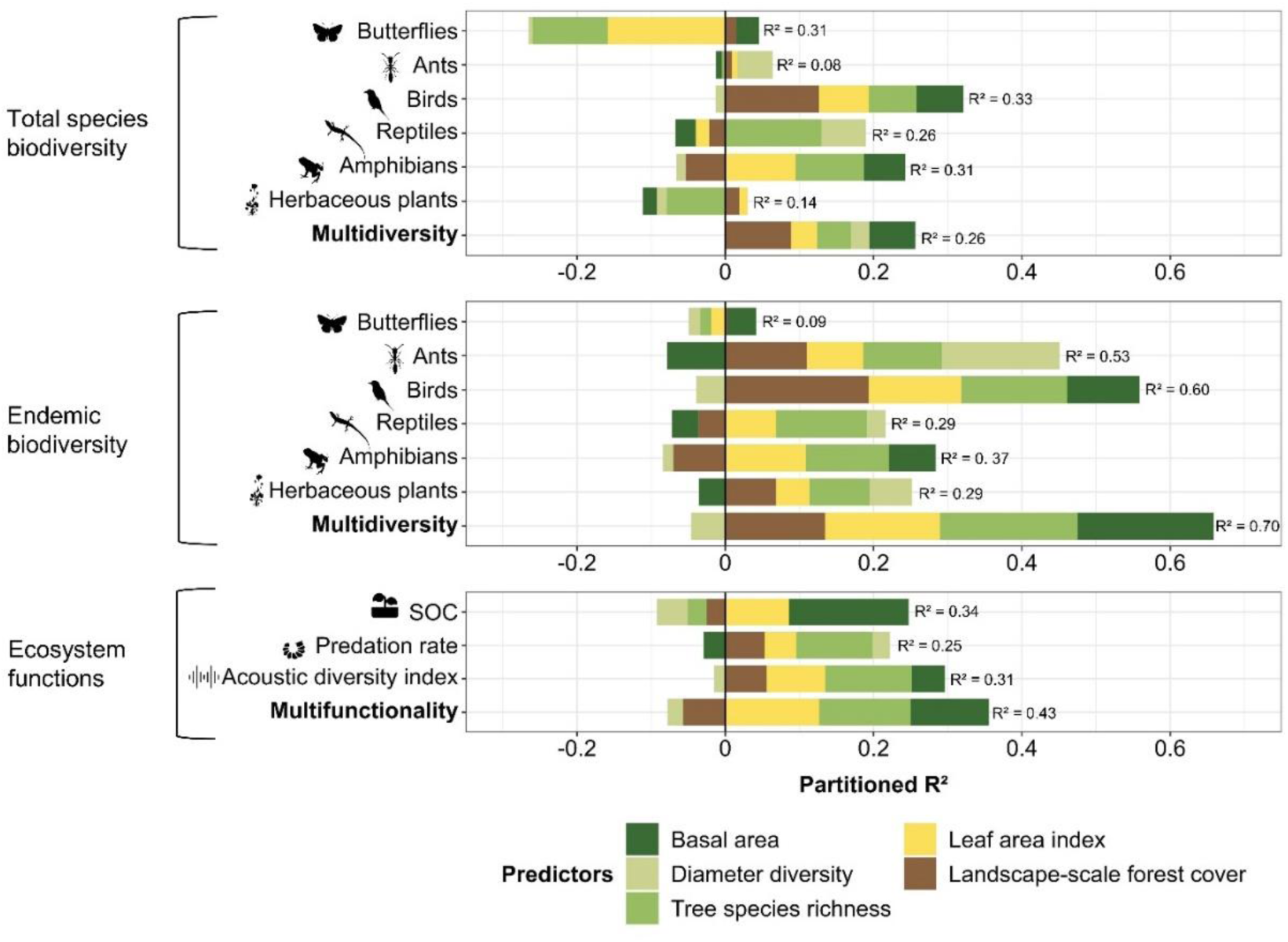
Relative importance of plot-scale stand structure and forest landscape-scale cover for total species biodiversity, endemic biodiversity, and ecosystem functions in northeastern Madagascar based on hierarchical partitioning of R². Icons from phylopic.org and icons8.de (see Table A7 for attribution).

## 4. Discussion

Our results supported our hypothesis that a more complex stand structure and a higher landscape-scale forest cover are associated with higher species richness for endemic biodiversity (R² = 0.70) but lower for total species biodiversity (R² = 0.26). Similarly, they generally supported ecosystem functions (R² = 0.43). The explanatory power of each predictor varied among taxa and ecosystem functions. Total species richness responded negatively and positively, and endemic species richness responded positively to our predictors. Basal area and tree species richness were the most predictive variables for endemic multidiversity. Landscape-scale forest cover also played an important role in predicting the species richness of birds (total and endemic species).

### 4.1. Complex stand structure is most important for endemics

Models of endemic multidiversity accounted for 70% of the variation. Notably, larger basal area and higher tree species richness emerged as important predictors of endemic multidiversity. Our findings are in line with a study by Waltert et al. (2011), which showed how a reduction in habitat complexity resulted in a decline in the species richness of endemics (birds, butterflies, and understory plants) in Cameroonian forests. Our results indicate that many endemic species are forest-dependent (Goodman and Benstead, 2005). For instance, 13 of the 31 endemic bird species inventoried in the study region occurred solely in old-growth forests (Martin et al., 2021a). We assume that numerous endemic species are specialized for forest habitats and rely on habitat structural characteristics. In fact, a stand with a higher basal area implies higher biomass (Soazafy et al., 2021). The identity and composition of tree species within the diverse pool might hold particular relevance for forest-specialist species. Thus, stands with complex structure emerge as resource-rich habitats, benefiting endemic species. Importantly, transforming landscapes into simplified agricultural landscapes with fewer trees and less diverse stands would be at a high cost to endemic species. Our findings provide comprehensive evidence of the importance of conserving and restoring complex stand structure for endemic communities.

### 4.2. Landscape-scale forest cover drives bird species richness

Landscape-scale forest cover was particularly predictive and explained a large proportion of the total variance in biodiversity. Among taxa, birds responded strongly and positively to landscape-scale forest cover. This finding is consistent with a study by Kupsch et al. (2019) in southwestern Cameroon. Forest cover in the landscape is important for birds, which are highly mobile animals and require more space as a habitat, and they are sensitive to habitat fragmentation (Bregman et al., 2014). Maintaining a certain threshold of forest cover in the landscapes plays an important role for forest species, which require certain areas as habitats (Arroyo-Rodríguez et al., 2020). Our results imply that conservation and restoration actions are not restricted to a limited area but should be planned and implemented on a landscape scale (Von Holle et al., 2020).

### 4.3. Stand structure is less predictive of total species richness

Stand structure and landscape-scale forest cover weakly predicted multidiversity. This aligns with the findings of Ampoorter et al. (2020), indicating that forest structure had a limited effect on multidiversity. Responses of each taxon to our predictors differed, resulting in a less pronounced effect on the combined metric, multidiversity. Indeed, each species/taxon has its own habitat requirements. We found that low LAI was correlated with high butterfly species richness, likely because light provides favorable conditions for nectar collection, basking, mating, and other activities butterflies love to engage in (Vlasanek et al., 2013). Conversely, high LAI was also correlated with high amphibian species richness, presumably due to a more humid microclimate suitable for tropical amphibian forest species (Veselka et al., 2023). The use of combined metrics such as multidiversity is useful to deal with these different responses. Moreover, changes in stand structure characteristics may have a lesser effect on total species richness, given that forest conversion favored certain exotic or native species (Rakotomalala et al., 2021; Raveloaritiana et al., 2021). Our results highlight the importance of using endemic species as a biodiversity metric rather than total species, particularly in modified habitats (De Lima et al., 2013; Waltert et al., 2011).

### 4.4. Difference in response among taxa

Total species richness responded negatively (butterflies, herbaceous plants) and positively (other taxa) to plot-scale stand structure and landscape-scale forest cover, whereas endemic species richness generally responded positively. These results reiterate that endemic taxa would profit more from complex stand structure. In addition, they suggest that some species of butterflies and herbaceous plants would prefer open areas (Schultz et al., 2012), such as grasslands. In biodiversity conservation efforts, it is essential to consider the distinct habitat requirements of target taxa, which can vary between different groups.

### 4.5. Stand structure as a driver of ecosystem functions

Basal area drives SOC stock. Hombegowda et al. (2016) reported similar results in southern India. Big trees provide a high quantity of litter and root biomass, which are carbon inputs in soil (Hu et al., 2016). Tree species richness was also significant in predicting acoustic diversity. The result is concordant with Beason et al. (2023) in temperate forests. Apparently, multiple tree species provide a habitat for biodiversity, which manifests itself in a variety of vocalizations. In fact, acoustic diversity in the study region was positively associated with bird species richness (Dröge et al., 2021). For multifunctionality, our models explained 43% of the variance, but there were no significant predictors. The relationships manifest themselves as a correlation between multifunctionality and basal area, tree species richness, and LAI. In fact, greater tree species richness increased litter diversity, affecting litter decomposition (Beugnon et al., 2023), and improved resistance to pests by reducing host trees (Jactel et al., 2021). LAI is an indicator of a microclimate with cooler temperatures and higher humidity in tropical forests (Hardwick et al., 2015). Microclimates are recognized as having a strong influence on forest functioning through their effect on soil dynamics (De Frenne et al., 2021). The findings reported here may provide some early insights into the relationship between stand structure and ecosystem functions, but more comprehensive studies are needed with more measures of ecosystem functions.

### 4.6. Conservation and restoration implications

Our study focused on direct drivers of biodiversity and ecosystem functions. Land-use change, forest use, forest management, agroforestry management, and ecosystem restoration all influence stand structure, which then - as demonstrated here - drives biodiversity and ecosystem functions. Complex plot-scale stand structure, such as higher basal area and greater tree species richness, should be maintained in forests and can serve as target variables in restoration activities. The study region, where the original tropical moist forests have been converted into agricultural mosaic landscapes, can benefit from this knowledge. Vanilla agroforests can contribute to the preservation of endemic biodiversity and ecosystem functions by conserving complex stand structure in forest-derived agroforests and restoring it in fallow-derived agroforests. Furthermore, the remaining forest cover in the study region is important for (endemic) biodiversity (Martin et al., 2021a; Rakotomalala et al., 2021), and outcomes of reforestation projects (MEDD and IUCN, 2020) or biodiversity-friendly agricultural practices remain difficult to predict (Martin et al., 2021b). Probably, the rapid and ongoing forest loss in this initially forested landscape (Vieilledent et al., 2018) has already led to a decline in biodiversity and ecosystem functioning to a state from which a recovery can take a hysteretic and slow pathway. Consequently, restoration efforts might not lead to initial biodiversity levels, and preservation of remaining forests hence remains essential to protecting endemic biodiversity and associated ecosystem functions.

## Ethics statement

The fieldwork was conducted under research permits N°100/17/MEEF/SG/DGF/DSAP/SCB.Re, N°163/17/MEEF/SG/DGF/DSAP/SCB.Re, N°18/18/ MEEF/SG/DGF/DSAP/SCB.Re, and N°254/18/MEEF/ SG/DGF/DSAP/SCB.Re.

## CRediT authorship contribution statement

Ri.R: conceptualization, formal analysis, methodology, visualization, writing-original draft, writing-review and editing; Y.K.: conceptualization, methodology, supervision, writing-review and editing; S.D., T.R.F., K.O., A.A.N.A.R., Ro.R., E.R., D.S., M.R.S., and A.W.: investigation, writing-review and editing; I.G., H.K., B.R., H.L.T.R., F.M.R., and T.T.: supervision, writing-review and editing; A.A. and L.H.R.R: supervision; D.H. and D.A.M: conceptualization, methodology, supervision. writing-review and editing.

All authors contributed to the article and approved the submitted version.

## Acknowledgments

We thank all research assistants participating in the field data collection and all plot owners for access to their land.

We thank the German Academic Exchange Service (DAAD) under the “Development-Related Postgraduate Courses (EPOS)” program.

## Appendix A

Supplementary material

## Data Availability

Data and R-scripts are available on the OSF platform under https://osf.io/mqwd7/?view_only=fa7ebe9ed42f448295c492271d8345a8

