## Supplementary material for "Stand structure as the proximate driver of endemic biodiversity and ecosystem functions in tropical mosaic landscapes": Supplementary material_bioRxiv.docx

**Rajaonarimalala et al.**

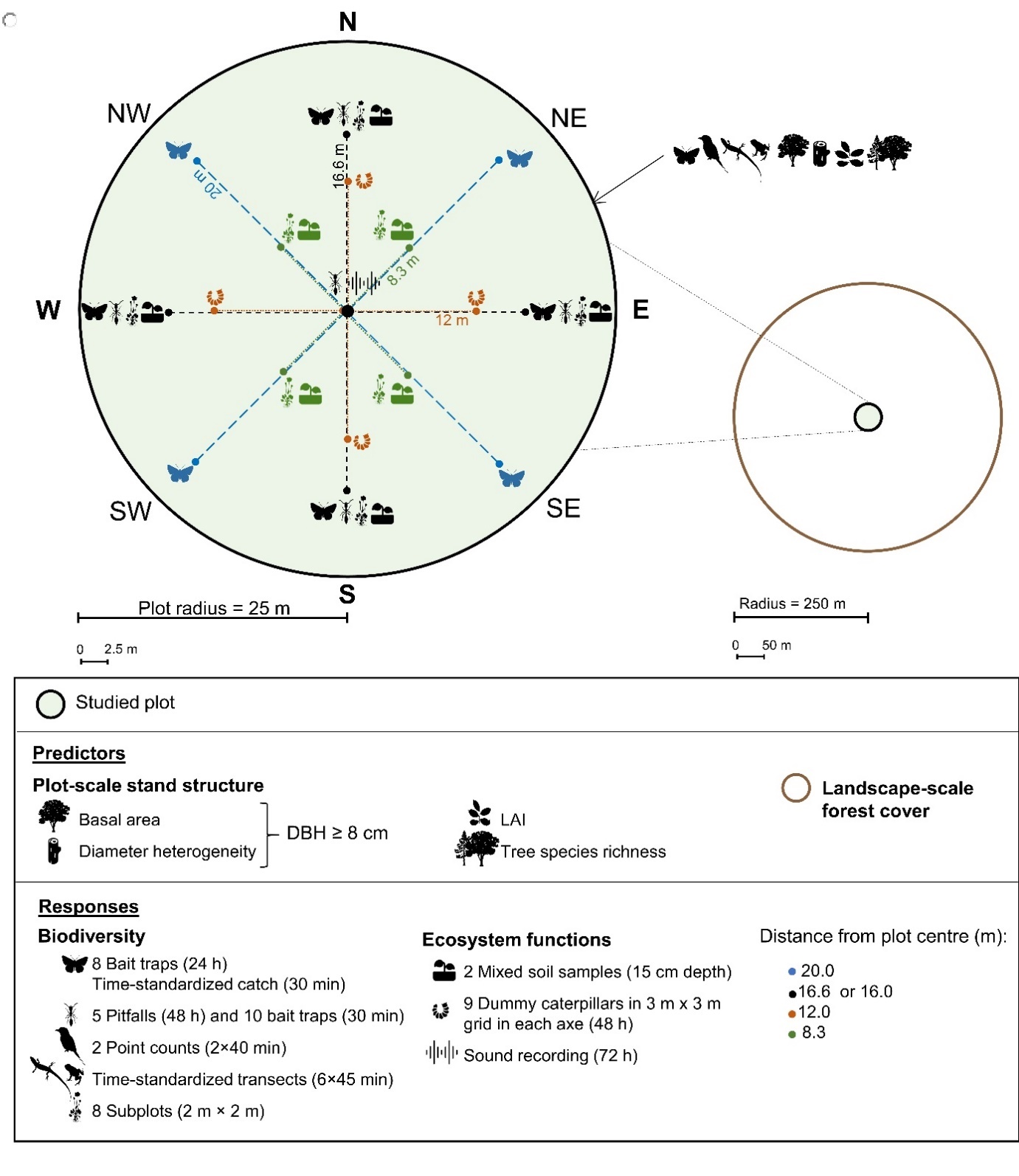

Fig. A1. Data collection design.

We collected data of **s**tand structural attributes within 25 m radius the plot (Osen et al., 2021; Soazafy et al., 2021), and estimated landscape-scale forest cover within a 250 m radius around the plot center (Martin et al., 2021). We caught or counted butterflies, birds, reptiles and amphibians inside the whole plot. We collected other taxa and ecosystem function variables at different locations within the plot (Dröge et al., 2021; Fulgence et al., 2022; Martin et al., 2022, 2021; Rakotomalala et al., 2021; Raveloaritiana et al., 2021; Schwab et al., 2021; Wurz et al., 2022). The dashed lines represent distances from the plot center. Icons from phylopic.org and icons8.de (see table S7 for attribution).

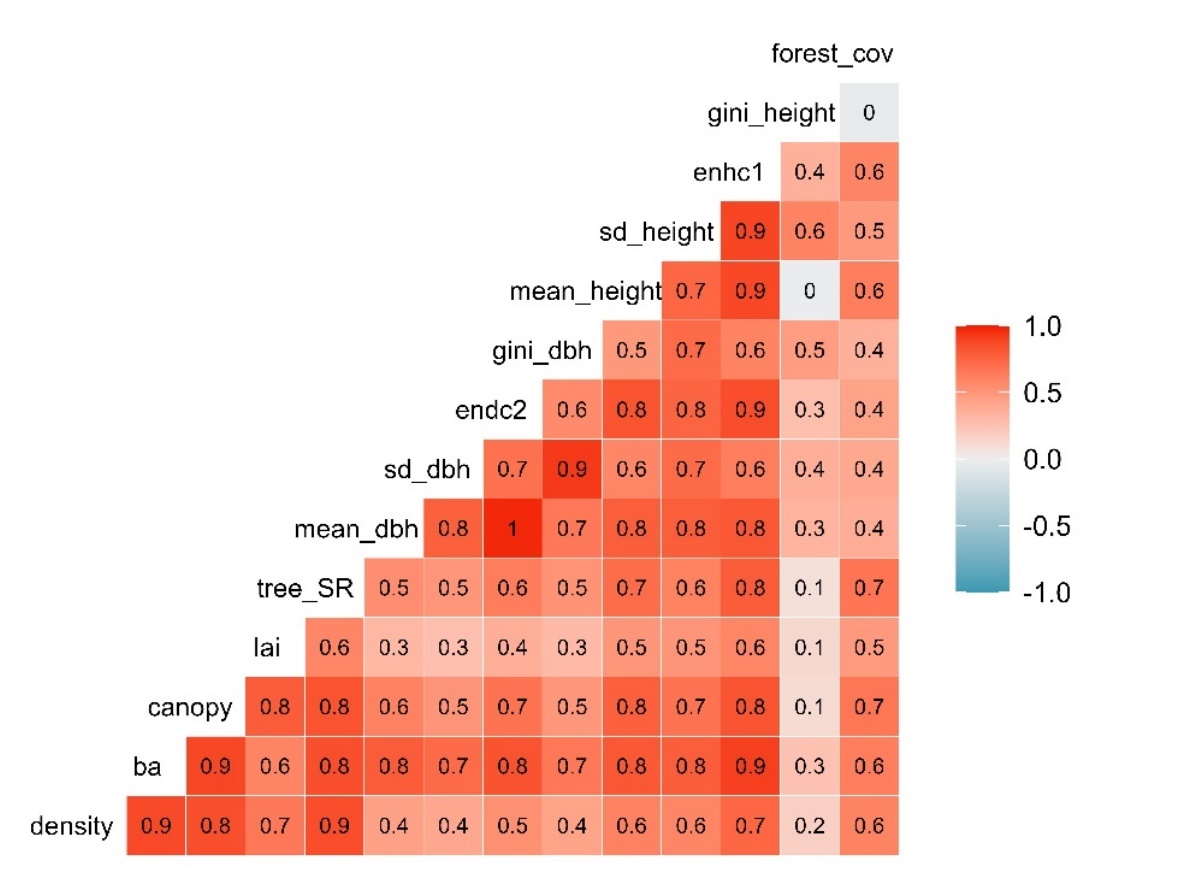

Fig. A2. Pairwise Spearman correlation between predictors

: plot-scale stand structure and landscape-scale forest cover (R package *GGally* (Schloerke et al., 2023)). **density**: stem density (n stems per ha), **ba**: basal area (m^2^/ha), **canopy:** canopy closure (%), **lai**: leaf area index (m^2^ _leaf area_/m^2^ _ground_), t**ree_SR**: tree species richness (number of tree species per plot), **mean_dbh**: mean diameter (cm), **sd_dbh**:  diameter heterogeneity: standard deviation (cm), **endc2**: diameter diversity: effective number of tree DBH classes (width of the class = 2 cm), **gini_dbh**: diameter inequality: Gini coefficient of tree diameter, **mean_height**: mean height (m), **sd_height**: height heterogeneity: standard deviation (m), **enhc_1**: height diversity: effective number of tree height classes (width of the class = 1 m), **gini_height**:  height inequality: Gini coefficient of tree height., **forest_cov:** proportion of forest area within 250 m radius around plot center (%).

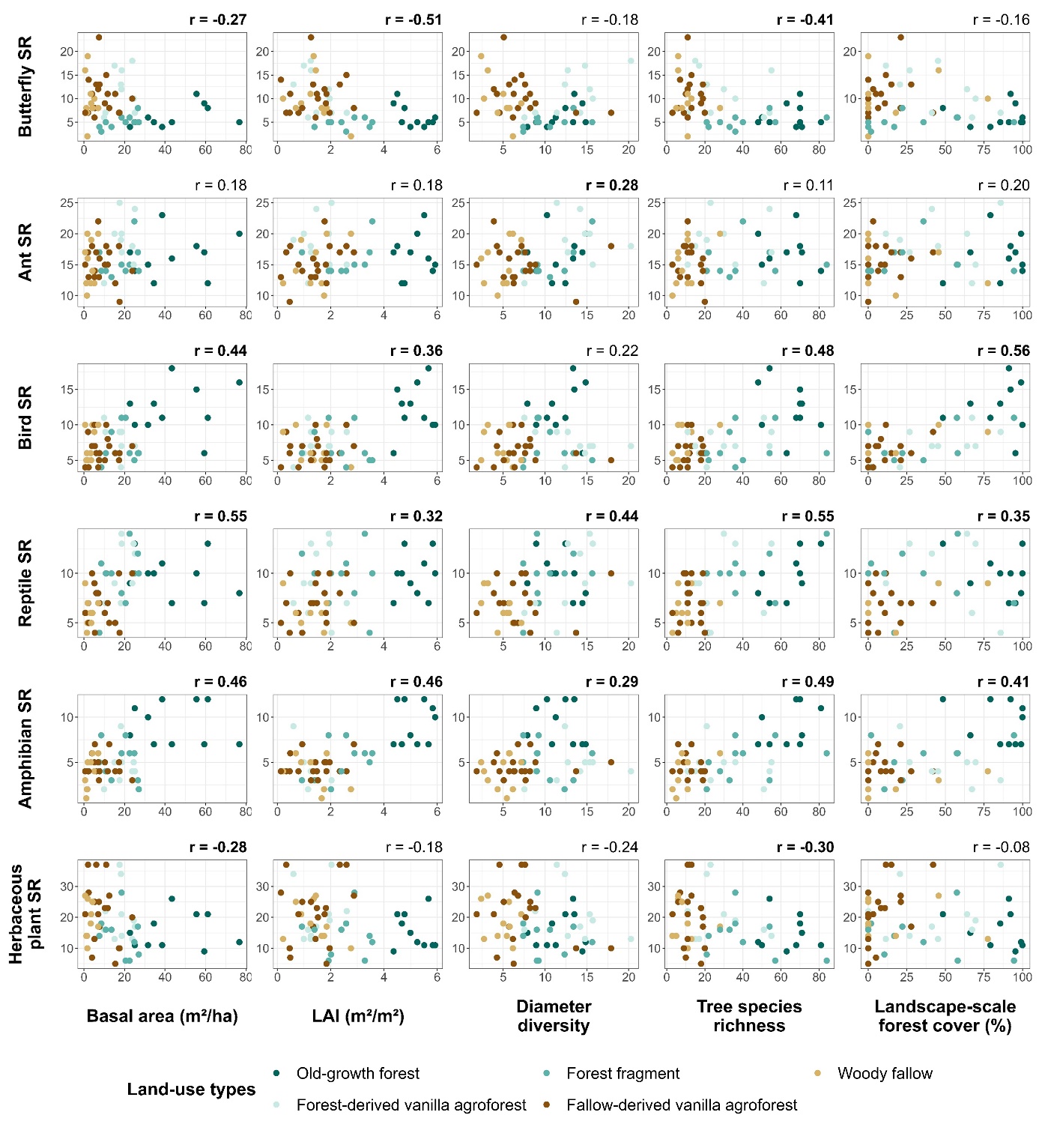

Fig. A3. Correlation analyses between predictors and total species richness.

Predictors are four plot-scale stand structure: basal area, leaf area index (LAI), diameter diversity, and tree species richness; and also, landscape-scale forest cover. Response variables are total species richness (SR) of six studied taxa. Scatterplots are colored by land-use type. r is the Spearman correlation coefficient and significant correlation is in bold font (p < 0.05).

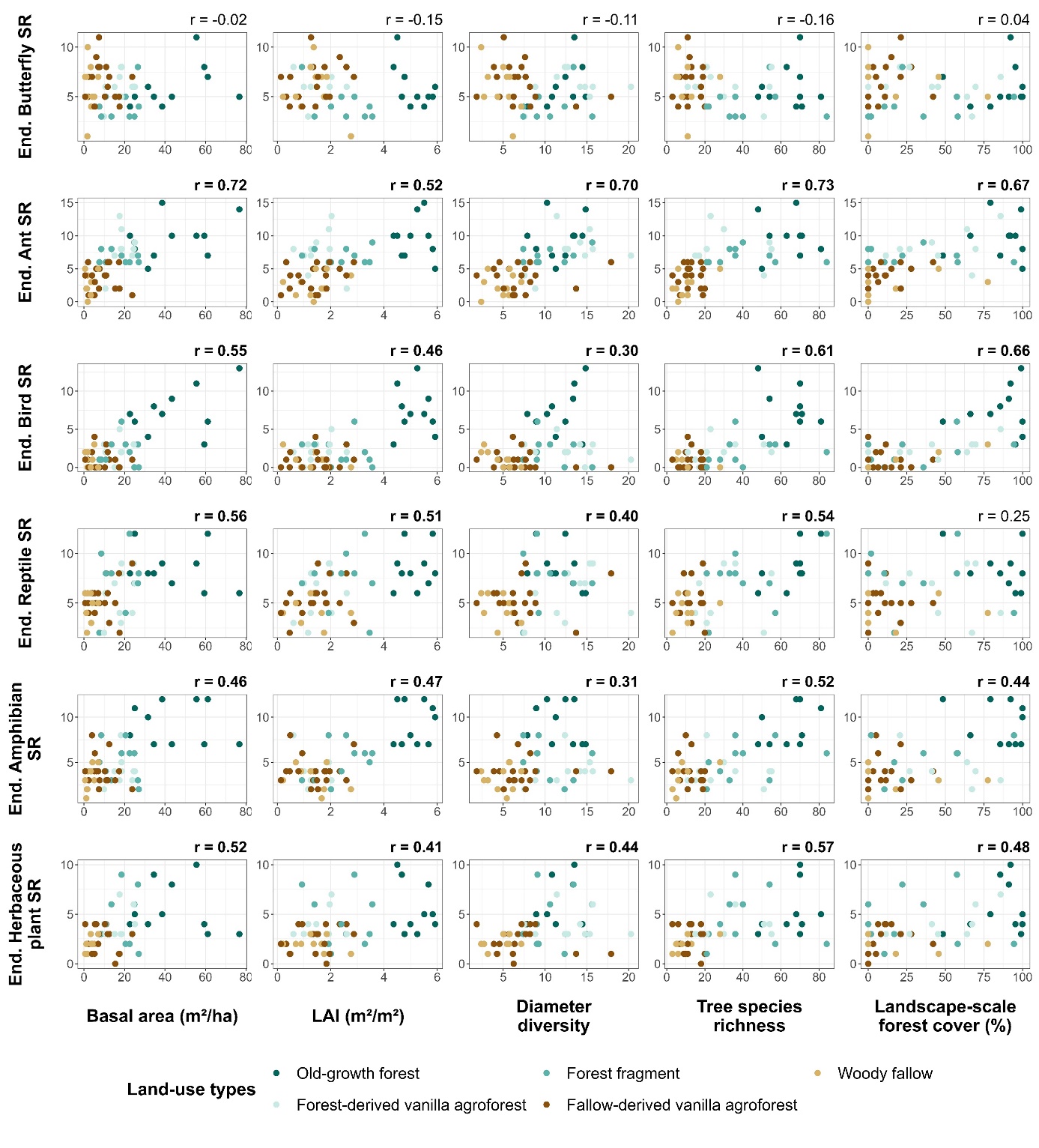

Fig. A4. Correlation analyses between predictors and endemic species richness.

Predictors are four plot-scale stand structure: basal area, leaf area index (LAI), diameter diversity, and tree species richness; and also, landscape-scale forest cover. Response variables are endemic (End.) species richness (SR) of six studied taxa. Scatterplots are colored by land-use type. r is the Spearman correlation coefficient and significant correlation is in bold font (p < 0.05).

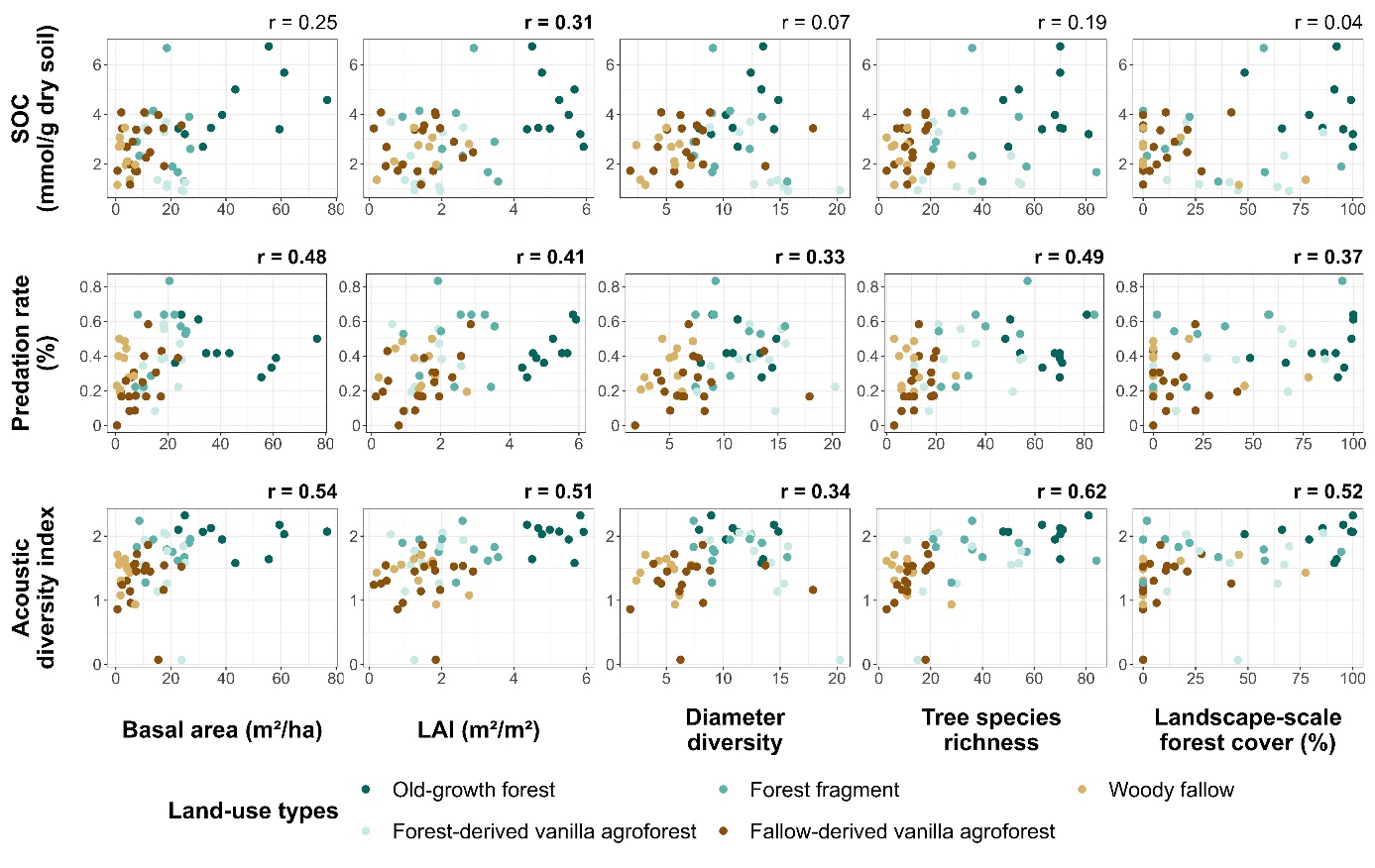

Fig. A5. Correlation analyses between predictors and ecosystem functions.

Predictors are four plot-scale stand structure: basal area, leaf area index (LAI), diameter diversity, and tree species richness; and also, landscape-scale forest cover. Response variables are three ecosystem functions: soil organic carbon (SOC), predation rate, and acoustic diversity index. Scatterplots are colored by land-use type. r is the Spearman correlation coefficient and significant correlation is in bold font (p < 0.05).

Table A1. Method description of some plot-scale stand structure.

| **Variable** | **Method** |
| --- | --- |
| Mean tree diameter (cm) and mean tree height (m).  Stem density (number of stems per ha). | Tree inventory all trees with diameter at breast height greater than 8cm. A vertex III (Haglöf, Långsele, Sweden) to evaluate tree height (Osen et al., 2021; Soazafy et al., 2021). |
| Standard deviation of diameter (cm) and height (m) measurements. | $SD=exp(-\sum_{i=1}^{n} {(x_{i}-x̄)}^{2})$ (A.1)  $x_{i}$: Tree diameter or tree height measurement  $x̄$: Mean of tree diameter or tree height  n: Number of measured trees |
| Diameter and height diversity:  Effective number of DBH classes (ENDC; Ehbrecht et al., 2019) of 2 cm each and height classes (ENHC) of 1 m each, were computed using package *hillR* in R (Li, 2018). | $ENDC or ENHC=exp(-\sum_{i=1}^{S} p_{i}ln p_{i})$ (A.2)  $p_{i}$: Proportion of trees in $i^{th}$ tree diameter or tree height class  $S$. Total number of tree diameter or tree height classes in each plot  (Jost, 2006) |
| Gini coefficients of diameter (GC_d_) and height (GC_h_) were calculated with R package *ineq* (Zeileis, 2014), to evaluate the diameter and height inequality of the trees. | ${GC}_{d}or {GC}_{h}=(\sum_{i=1}^{n} \left( 2i-n-1 \right)x_{i})/( n\sum_{i=1}^{n} x_{i})$ (A.3)  $x$: Tree diameter or tree height measurement  $i$: Rank of the values  $n$: Number of tree diameter or tree height measurement in each plot  (Gini, 1912) |
| Canopy closure (%). | Nikon D5100 camera (Nikon, Tokyo, Japan) to take canopy pictures.  Canopy closure percentage obtained with “ImageJ” software (Rasband, 2014) separating vegetation from the sky through binary image classification (Osen et al., 2021). |

Table A2. Model assessment of GLM results.

Functions used: (1) check_collinearity(), (2) testUniformity(), (3) check_overdispersion(), (4) testDispersion(), (5) testQuantiles(), (6) testZeroInflation(), (7) check_singularity(), (8) check_convergence() with R packages *performance* (Lüdecke et al., 2021) and *DHARMa* (Hartig, 2022). VIF: variance inflation factor, SR: species richness, End.: endemic, SOC: soil organic carbon, ba: basal area, lai: leaf area index, endc2: diameter diversity, tree_SR: tree species richness, forest_cov: landscape-scale forest cover.

| **Model** | **Family** | **Collinearity**  **VIF values** | **Uniformity of residuals** | **Over-**  **dispersion** | **Dispersion of residuals** | **Heterosce-dasticity of residuals** | **Zero inflation** | **Singularity** | **Convergence** |
| --- | --- | --- | --- | --- | --- | --- | --- | --- | --- |
|  |  | (1) | (2) | (3) | (4) | (5) | (6) | (7) | (8) |
| Butterfly SR | Negative binomial | Low:  ba: 4.01, lai: 2.34, endc2: 1.90  tree SR: 2.68, forest_cov: 1.97 | p-value = 0.8742 | p-value = 0.180 | p-value = 0.48 | p-value = 0.8917 | p-value = 1 | No | No |
| Ant SR | Poisson | Low:  ba: 3.87 , lai: 2.64 , endc2: 1.84  tree SR: 2.90, forest_cov: 2.41 | p-value = 0.1519 | p-value = 0.951 | p-value = 0.016 (a) | p-value = 0.992 | p-value = 1 | No | No |
| Bird SR | Poisson | Low:  ba: 3.90 , lai: 3.00 , endc2: 1.97  tree SR: 2.99, forest_cov: 2.54 | p-value = 0.5294 | p-value = 0.975 | p-value = 0.024 (a) | p-value = 0.9405 | p-value = 1 | No | No |
| Reptile SR | Poisson | Low:  ba: 3.72, lai: 2.71 , endc2: 1.83  tree SR: 2.75, forest_cov: 2.52 | p-value = 0.5777 | p-value = 0.991 | p-value = 0.008 (a) | p-value = 0.6401 | p-value = 1 | No | No |
| Amphibian SR | Poisson | Low:  ba: 3.75, lai: 3.08 , endc2: 1.93  tree SR: 2.97, forest_cov: 2.84 | p-value = 0.3436 | p-value = 0.98 | p-value = 0.04 (a) | p-value = 0.026 (b) | p-value = 1 | No | No |
| Herbaceous plant SR | Negative binomial | Low:  ba: 3.85, lai: 2.43 , endc2: 1.91  tree SR: 2.93, forest_cov: 2.29 | p-value = 0.7135 | p-value = 0.315 | p-value = 0.936 | p-value = 0.9004 | p-value = 1 | No | No |
| Multi-diversity | Gaussian | Low:  ba: 3.92 , lai: 2.61 , endc2: 1.90  tree SR: 2.93, forest_cov: 2.37 | p-value = 0.7949 | - | - | p-value = 0.9941 | - | No | No |
| End. Butterfly SR | Poisson | Low:  ba: 4.12 , lai: 2.74 , endc2: 1.96  tree SR: 2.94, forest_cov: 2.30 | p-value = 0.4425 | p-value = 0.976 | p-value = 0.04 (a) | p-value = 0.7798 | p-value = 1 | No | No |
| End. Ant SR | Poisson | Low:  ba: 3.48, lai: 3.00 , endc2: 1.68  tree SR: 2.65, forest_cov: 2.66 | p-value = 0.9484 | p-value = 0.575 | p-value = 0.576 | p-value = 0.3888 | p-value = 1 | No | No |
| End. Bird SR | Poisson | Low:  ba: 3.72 , lai: 3.20 , endc2: 2.32  tree SR: 2.49, forest_cov: 2.52 | p-value = 0.9897 | p-value = 0.271 | p-value = 0.99 | p-value = 0.3438 | p-value = 0.68 | No | No |
| End. Reptile SR | Poisson | Low:  ba: 3.68, lai: 2.90 , endc2: 1.85  tree SR: 2.89, forest_cov: 2.81 | p-value = 0.3309 | p-value = 0.993 | p-value < 0.001 (a) | p-value = 0.644 | p-value = 1 | No | No |
| End. Amphibian SR | Poisson | Low:  ba: 3.71 , lai: 3.14 , endc2: 1.96  tree SR: 2.96, forest_cov: 2.88 | p-value = 0.5535 | p-value = 0.944 | p-value = 0.064 (a) | p-value = 0.004 (b) | p-value = 0.768 | No | No |
| End. Herbaceous plant SR | Poisson | Low:  ba: 3.58 , lai: 2.89 , endc2: 1.77  tree SR: 2.75, forest_cov: 2.61 | p-value = 0.78 | p-value = 0.628 | p-value = 0.856 | p-value = 0.9411 | p-value = 0.328 | No | No |
| End. Multidiversity | Gaussian | Low:  ba: 3.92 , lai: 2.61 , endc2: 1.90  tree SR: 2.93, forest_cov: 2.37 | p-value = 0.6377 | - | - | p-value = 0.8365 | - | No | No |
| SOC | Gaussian | Low:  ba: 3.92 , lai: 2.61 , endc2: 1.90  tree SR: 2.93, forest_cov: 2.37 | p-value = 0.5884 | - | - | p-value = 0.7053 | - | No | No |
| Predation | Gaussian | Low:  ba: 3.92 , lai: 2.61 , endc2: 1.90  tree SR: 2.93, forest_cov: 2.37 | p-value = 0.2141 | - | - | p-value = 0.746 | - | No | No |
| Acoustic diversity index | Gaussian | Low:  ba: 3.92 , lai: 2.61 , endc2: 1.90  tree SR: 2.93, forest_cov: 2.37 | p-value = 0.3599 |  |  | p-value = 0.2273 | - | No | No |
| Multi-functionality | Gaussian | Low:  ba: 3.92 , lai: 2.61 , endc2: 1.90  tree SR: 2.93, forest_cov: 2.37 | p-value = 0.6501 | - | - | p-value = 0.8267 | - | No | No |

(Note: (4; a) Cases of underdispersion were addressed using the Conway Maxwell-Poisson distribution (Brooks et al., 2017). However, R² could not be obtained for this distribution, that gave residual variance instead (Bartoń, 2023), so Poisson distribution was used. In fact, coefficient estimates and AIC of Conway Maxwell-Poisson and Poisson model results were almost the same; (5; b) Cases of heteroscedasticity of residuals; ().

Table A3. Correlation matrix between predictors and responses.

Correlation using Spearman method, correlation coefficient ≥ 0.50 in grey and significant correlation in bold font (p < 0.05), density: stem density (n stems per ha), ba: basal area (m²/ha), canopy: canopy closure (%), lai: leaf area index (m² leaf area/m² ground), tree_SR: tree species richness (number of tree species per plot), mean_dbh: mean diameter (cm), sd_dbh: diameter heterogeneity: standard deviation (cm), endc2: diameter diversity: effective number of tree DBH classes (width of the class = 2 cm), gini_dbh: diameter inequality: Gini coefficient of tree diameter, mean_height: mean height (m), sd_height: height heterogeneity: standard deviation (m), enhc_1: height diversity, effective number of tree height classes (width of the class = 1 m), gini_height: height inequality: Gini coefficient of tree height, forest_cov: Proportion of forest area within 250 m radius around plot center (%). SR: species richness, End.: endemic, SOC: soil organic carbon.

| **Predictors** | density | ba | canopy | lai | tree_  SR | mean_  dbh | sd_  dbh | endc2 | gini_  dbh | mean_  height | sd_  height | enhc1 | gini_  height | forest_cov |
| --- | --- | --- | --- | --- | --- | --- | --- | --- | --- | --- | --- | --- | --- | --- |
| **Responses** |  |  |  |  |  |  |  |  |  |  |  |  |  |  |
| Butterfly SR | **-0.31** | **-0.27** | **-0.39** | **-0.51** | **-0.41** | -0.10 | -0.07 | -0.18 | -0.19 | **-0.27** | -0.25 | **-0.34** | -0.06 | -0.16 |
| Ant SR | 0.09 | 0.18 | **0.28** | 0.18 | 0.11 | **0.27** | 0.12 | **0.28** | 0.02 | **0.29** | 0.19 | 0.25 | 0.02 | 0.20 |
| Bird SR | **0.46** | **0.44** | **0.41** | **0.36** | **0.48** | 0.21 | **0.30** | 0.22 | **0.33** | **0.36** | **0.39** | **0.42** | 0.07 | **0.56** |
| Reptile SR | **0.42** | **0.55** | **0.51** | **0.32** | **0.55** | **0.43** | **0.38** | **0.44** | 0.25 | **0.50** | **0.42** | **0.47** | -0.05 | **0.35** |
| Amphibian SR | **0.48** | **0.46** | **0.54** | **0.46** | **0.49** | 0.25 | **0.30** | **0.29** | **0.31** | **0.46** | **0.39** | **0.44** | -0.01 | **0.41** |
| Herbaceous plant SR | -0.21 | **-0.28** | -0.19 | -0.18 | **-0.30** | -0.18 | -0.26 | -0.24 | -0.20 | **-0.31** | -0.07 | -0.21 | 0.23 | -0.08 |
| Multidiversity | **0.38** | **0.43** | **0.47** | **0.31** | **0.37** | **0.33** | **0.26** | **0.32** | 0.18 | **0.40** | **0.39** | **0.41** | 0.08 | **0.54** |
| End. Butterfly SR | 0.04 | -0.02 | -0.04 | -0.15 | -0.16 | -0.08 | -0.03 | -0.11 | -0.08 | -0.13 | -0.10 | -0.13 | 0.01 | 0.04 |
| End. Ant SR | **0.59** | **0.72** | **0.76** | **0.52** | **0.73** | **0.63** | **0.46** | **0.70** | **0.40** | **0.76** | **0.63** | **0.75** | 0.10 | **0.67** |
| End. Bird SR | **0.55** | **0.55** | **0.60** | **0.46** | **0.61** | **0.28** | **0.38** | **0.30** | **0.42** | **0.49** | **0.48** | **0.55** | 0.07 | **0.66** |
| End. Reptile SR | **0.53** | **0.56** | **0.56** | **0.51** | **0.54** | **0.38** | **0.29** | **0.40** | 0.20 | **0.46** | **0.41** | **0.48** | -0.04 | 0.25 |
| End. Amphibian SR | **0.47** | **0.46** | **0.56** | **0.47** | **0.52** | 0.25 | **0.27** | **0.31** | **0.30** | **0.47** | **0.39** | **0.47** | -0.01 | **0.44** |
| End. Herbaceous plant SR | **0.52** | **0.52** | **0.53** | **0.41** | **0.57** | **0.41** | **0.30** | **0.44** | **0.33** | **0.47** | **0.58** | **0.61** | **0.27** | **0.48** |
| End. Multidiversity | **0.72** | **0.74** | **0.79** | **0.60** | **0.74** | **0.49** | **0.42** | **0.54** | **0.40** | **0.65** | **0.61** | **0.70** | 0.11 | **0.65** |
| SOC | **0.38** | 0.25 | **0.27** | **0.31** | 0.19 | 0.03 | 0.06 | 0.07 | 0.15 | 0.04 | 0.23 | 0.23 | **0.37** | 0.04 |
| Predation rate | **0.48** | **0.48** | **0.57** | **0.41** | **0.49** | **0.26** | 0.21 | **0.33** | 0.19 | **0.39** | **0.27** | **0.40** | -0.08 | **0.37** |
| Acoustic diversity index | **0.61** | **0.54** | **0.64** | **0.51** | **0.62** | **0.26** | 0.22 | **0.34** | **0.30** | **0.50** | **0.44** | **0.56** | 0.04 | **0.52** |
| Multifunctionality | **0.67** | **0.59** | **0.71** | **0.57** | **0.60** | **0.29** | 0.23 | **0.37** | **0.32** | **0.46** | **0.46** | **0.56** | 0.14 | **0.45** |

Table A4. Effect of plot-scale stand structure and landscape-scale forest cover on (endemic) biodiversity (six taxa), and ecosystem functions (three functions) from a GLM.

Significant predictors (p < 0.05) in bold font. GLM: Generalized Linear Model. Est.: estimate, SE: standard error, R²: coefficient of determination, End.: endemic, LAI: leaf area index, SR: species richness, SOC: soil organic carbon.

| Responses |  | Intercept | Basal area | LAI | Diameter diversity | Tree SR | Landscape-scale forest cover | R^2^ |
| --- | --- | --- | --- | --- | --- | --- | --- | --- |
| **Total species biodiversity** | | | | | | | | |
| Butterfly SR | Est. | 2.15 | 0.16 | **-0.26** | -0.01 | -0.19 | 0.08 | 0.31 |
|  | SE | 0.05 | 0.11 | **0.09** | 0.07 | 0.10 | 0.08 |  |
|  | z value | 39.49 | 1.46 | **-2.89** | -0.10 | -1.96 | 1.00 |  |
| Ant SR | Est. | 2.77 | -0.03 | 0.04 | 0.08 | -0.04 | 0.03 | 0.08 |
|  | SE | 0.03 | 0.06 | 0.05 | 0.04 | 0.06 | 0.05 |  |
|  | z value | 84.48 | -0.53 | 0.75 | 1.75 | -0.76 | 0.68 |  |
| Bird SR | Est. | 2.02 | 0.11 | 0.03 | -0.09 | 0.01 | **0.17** | 0.33 |
|  | SE | 0.05 | 0.08 | 0.07 | 0.07 | 0.08 | **0.07** |  |
|  | z value | 41.79 | 1.32 | 0.44 | -1.30 | 0.09 | **2.48** |  |
| Reptile SR | Est. | 2.09 | -0.05 | -0.03 | 0.10 | **0.23** | -0.05 | 0.26 |
|  | SE | 0.05 | 0.09 | 0.07 | 0.06 | **0.07** | 0.07 |  |
|  | z value | 44.92 | -0.53 | -0.35 | 1.55 | **3.14** | -0.74 |  |
| Amphibian SR | Est. | 1.63 | 0.07 | 0.15 | -0.04 | 0.13 | -0.02 | 0.31 |
|  | SE | 0.06 | 0.10 | 0.09 | 0.08 | 0.09 | 0.09 |  |
|  | z value | 27.68 | 0.76 | 1.64 | -0.47 | 1.47 | -0.19 |  |
| Herbaceous | Est. | 2.90 | -0.01 | 0.01 | -0.03 | **-0.20** | 0.11 | 0.14 |
| Plant SR | SE | 0.05 | 0.11 | 0.08 | 0.07 | **0.09** | 0.08 |  |
|  | z value | 54.04 | -0.12 | 0.11 | -0.50 | **-2.14** | 1.33 |  |
| **Endemic biodiversity** | | | | | | | | |
| End. Butterfly | Est. | 1.75 | **0.22** | -0.12 | -0.11 | -0.09 | 0.03 | 0.09 |
| SR | SE | 0.05 | **0.10** | 0.09 | 0.08 | 0.10 | 0.08 |  |
|  | z value | 31.98 | **2.17** | -1.36 | -1.44 | -0.94 | 0.34 |  |
| End. Ant SR | Est. | 1.66 | -0.04 | 0.07 | **0.25** | 0.10 | 0.16 | 0.53 |
|  | SE | 0.06 | 0.09 | 0.08 | **0.07** | 0.09 | 0.08 |  |
|  | z value | 27.64 | -0.41 | 0.86 | **3.50** | 1.23 | 1.92 |  |
| End. Bird SR | Est. | 0.55 | **0.32** | 0.05 | -0.23 | 0.16 | **0.45** | 0.60 |
|  | SE | 0.11 | **0.12** | 0.13 | 0.15 | 0.13 | **0.13** |  |
|  | z value | 4.97 | **2.72** | 0.40 | -1.61 | 1.27 | **3.50** |  |
| End. Reptile SR | Est. | 1.79 | <-0.01 | 0.12 | 0.05 | **0.24** | -0.17 | 0.29 |
|  | SE | 0.05 | 0.10 | 0.08 | 0.08 | **0.08** | 0.09 |  |
|  | z value | 32.97 | -0.01 | 1.36 | 0.63 | **2.87** | -1.95 |  |
| End. | Est. | 1.54 | 0.07 | 0.17 | -0.06 | 0.17 | <-0.01 | 0.37 |
| Amphibian SR | SE | 0.06 | 0.10 | 0.09 | 0.09 | 0.09 | 0.09 |  |
|  | z value | 24.78 | 0.77 | 1.79 | -0.64 | 1.79 | -0.02 |  |
| End. | Est. | 1.19 | -0.09 | 0.07 | 0.15 | 0.16 | 0.13 | 0.29 |
| Herbaceous | SE | 0.07 | 0.12 | 0.11 | 0.10 | 0.11 | 0.11 |  |
| Plant SR | z value | 15.98 | -0.70 | 0.62 | 1.57 | 1.45 | 1.22 |  |
| **Ecosystem functions** | | | | | | | | |
| SOC | Est. | 2.87 | **1.00** | 0.29 | **-0.49** | -0.09 | -0.35 | 0.34 |
|  | SE | 0.14 | **0.28** | 0.23 | **0.19** | 0.24 | 0.22 |  |
|  | z value | 20.51 | **3.58** | 1.26 | **-2.51** | -0.36 | -1.62 |  |
| Predation rate | Est. | 0.36 | -0.04 | 0.02 | 0.03 | 0.06 | 0.02 | 0.25 |
|  | SE | 0.02 | 0.04 | 0.03 | 0.03 | 0.03 | 0.03 |  |
|  | z value | 18.36 | -0.95 | 0.70 | 0.97 | 1.91 | 0.77 |  |
| Acoustic | Est. | 1.57 | 0.03 | 0.08 | -0.08 | **0.16** | 0.03 | 0.31 |
| diversity index | SE | 0.05 | 0.10 | 0.08 | 0.07 | **0.08** | 0.08 |  |
|  | z value | 32.52 | 0.31 | 0.98 | -1.12 | **1.97** | 0.42 |  |

Table A5. Model results: coefficient estimates in GLM results.

GLM: Response variable ~ basal area +LAI + diameter diversity + tree species richness + landscape-scale forest cover. GLM: Generalized Linear Model, AIC: Akaike information criterion, LAI: leaf area index, SR: species richness, End.: endemic, SOC: soil organic carbon.

| **Responses** | **Family** | **Estimates** | | | | | | **AIC** |
| --- | --- | --- | --- | --- | --- | --- | --- | --- |
|  |  | **Intercept** | **Basal area** | **LAI** | **Diameter diversity** | **Tree SR** | **Landscape-scale forest cover** |  |
| Butterfly SR | Negative binomial | 2.15 | 0.16 | -0.26 | -0.01 | -0.19 | 0.08 | 319.00 |
| Ant SR | Poisson | 2.77 | -0.03 | 0.04 | 0.08 | -0.04 | 0.03 | 314.90 |
| Bird SR | Poisson | 2.02 | 0.11 | 0.03 | -0.09 | 0.01 | 0.17 | 269.00 |
| Reptile SR | Poisson | 2.09 | -0.05 | -0.03 | 0.10 | 0.23 | -0.05 | 270.90 |
| Amphibian SR | Poisson | 1.63 | 0.07 | 0.15 | -0.04 | 0.13 | -0.02 | 245.80 |
| Herbaceous plant SR | Negative binomial | 2.90 | -0.01 | 0.01 | -0.03 | -0.20 | 0.11 | 406.90 |
| Multidiversity | Gaussian | 0.38 | 0.02 | <0.01 | 0.01 | <0.01 | 0.03 | -92.90 |
| End. Butterfly SR | Poisson | 1.75 | 0.22 | -0.12 | -0.11 | -0.09 | 0.03 | 253.70 |
| End. Ant SR | Poisson | 1.66 | -0.04 | 0.07 | 0.25 | 0.10 | 0.16 | 262.70 |
| End. Bird SR | Poisson | 0.55 | 0.32 | 0.05 | -0.23 | 0.16 | 0.45 | 197.30 |
| End. Reptile SR | Poisson | 1.79 | <0.01 | 0.12 | 0.05 | 0.24 | -0.17 | 253.90 |
| End. Amphibian SR | Poisson | 1.54 | 0.07 | 0.17 | -0.06 | 0.17 | <-0.01 | 241.70 |
| End. Herbaceous plant SR | Poisson | 1.19 | -0.09 | 0.07 | 0.15 | 0.16 | 0.13 | 231.80 |
| End. Multidiversity | Gaussian | 0.36 | 0.05 | 0.03 | <-0.01 | 0.04 | 0.02 | -112.20 |
| SOC | Gaussian | 2.87 | 1.00 | 0.29 | -0.49 | -0.09 | -0.35 | 185.90 |
| Predation | Gaussian | 0.36 | -0.04 | 0.02 | 0.03 | 0.06 | 0.02 | -41.90 |
| Acoustic diversity index | Gaussian | 1.57 | 0.03 | 0.08 | -0.08 | 0.16 | 0.03 | 62.80 |
| Multifunctionality | Gaussian | 0.48 | 0.05 | 0.04 | -0.03 | 0.05 | -0.01 | -74.3 |

Table A6. Model results: significance of GLM results.

GLM: Response variable ~ basal area +LAI + diameter diversity + tree species richness + forest cover. Significant p-value in bold font. SR: species richness. GLM: Generalized Linear Model, End.: endemic, SR: species richness, LAI: leaf area index, SOC: soil organic carbon.

| **Responses** | **Family** | **p-values** | | | | | |
| --- | --- | --- | --- | --- | --- | --- | --- |
|  |  | **Intercept** | **Basal area** | **LAI** | **Diameter diversity** | **Tree SR** | **Landscape-scale forest cover** |
| Butterfly SR | Negative binomial | **< 0.001** | 0.144 | **0.004** | 0.918 | 0.050 | 0.317 |
| Ant SR | Poisson | **< 0.001** | 0.593 | 0.451 | 0.080 | 0.450 | 0.493 |
| Bird SR | Poisson | **< 0.001** | 0.186 | 0.657 | 0.192 | 0.931 | **0.013** |
| Reptile SR | Poisson | **< 0.001** | 0.597 | 0.728 | 0.120 | **0.002** | 0.456 |
| Amphibian SR | Poisson | **< 0.001** | 0.446 | 0.102 | 0.642 | 0.142 | 0.850 |
| Herbaceous plant SR | Negative binomial | **< 0.001** | 0.908 | 0.909 | 0.618 | **0.032** | 0.183 |
| Multidiversity | Gaussian | **< 0.001** | 0.445 | 0.926 | 0.723 | 0.845 | 0.091 |
| End. Butterfly SR | Poisson | **< 0.001** | **0.030** | 0.173 | 0.149 | 0.349 | 0.734 |
| End. Ant SR | Poisson | **< 0.001** | 0.680 | 0.391 | <**0.001** | 0.217 | 0.055 |
| End. Bird SR | Poisson | **< 0.001** | **0.007** | 0.693 | 0.109 | 0.204 | <**0.001** |
| End. Reptile SR | Poisson | **< 0.001** | 0.991 | 0.172 | 0.527 | **0.004** | 0.051 |
| End. Amphibian SR | Poisson | **< 0.001** | 0.444 | 0.073 | 0.519 | 0.073 | 0.987 |
| End. Herbaceous plant SR | Poisson | **< 0.001** | 0.483 | 0.536 | 0.117 | 0.148 | 0.223 |
| End. Multidiversity | Gaussian | **< 0.001** | **0.021** | 0.095 | 0.815 | **0.016** | 0.198 |
| SOC | Gaussian | **< 0.001** | <**0.001** | 0.206 | **0.012** | 0.717 | 0.106 |
| Predation | Gaussian | **< 0.001** | 0.341 | 0.484 | 0.330 | 0.056 | 0.440 |
| Acoustic diversity index | Gaussian | **< 0.001** | 0.755 | 0.329 | 0.264 | **0.049** | 0.678 |
| Multifunctionality | Gaussian | **< 0.001** | 0.115 | 0.128 | 0.169 | 0.078 | 0.791 |

Table A7. Icon attribution.

| **Icon** | **Author** | **License** | **Link** |
| --- | --- | --- | --- |
| Butterfly | Matt Hayes | Public Domain | https://www.phylopic.org/images/ab6182d2-5093-444b-92e5-84468218ebf0/hamearis-lucina |
| Ant | Michael Day | Public Domain | https://www.phylopic.org/images/bfcdcccd-5ad9-44c5-b599-590d0422ee59/myrmica |
| Bird | Hugo Gruson | Public Domain | https://www.phylopic.org/images/086d2b94-f13c-47ab-9538-c9b71d97f934/coracias-caudata |
| Reptile | David Orr | Public Domain | https://www.phylopic.org/images/3dc70587-4e15-4f59-a246-4ec21390957b/plestiodon-fasciatus |
| Amphibian | Steven Traver | Public Domain | https://www.phylopic.org/images/4679516b-405b-444f-974d-9775876716e2/dendrobates-tinctorius |
| Herbaceous plant | Guillaume Dera | Public Domain | https://www.phylopic.org/images/6a5082eb-3162-414e-af1e-85bd3268ee68/parnassia-palustris |
| Soil organic carbon | Icons8 |  | https://icons8.de/icon/set/soil/group-filled |
| Predation rate | Icons8 |  | https://icons8.de/icons/set/larva |
| Acoustic diversity index | Icons8 |  | https://icons8.de/icons/set/wave |
| Basal area | Rachel T Mason | Public Domain | https://www.phylopic.org/images/3dc5c323-8094-4861-a853-3bb4de621c36/eucalyptus-obliqua |
| Diameter diversity | Icons8 |  | https://icons8.de/icon/9106/holz |
| Leaf area index | Ferran Sayol | Public Domain | https://www.phylopic.org/images/b5c81420-a49f-4d7a-8b82-1fcd94176889/hirtella-triandra |
| Tree species richness | Gabriele Midolo  Rachel T Mason | Public Domain | <https://www.phylopic.org/images/13a5a7e9-be33-4d40-8c90-9ffcf367d7cb/pinus-sect-trifoliae>  https://www.phylopic.org/images/3dc5c323-8094-4861-a853-3bb4de621c36/eucalyptus-obliqua |
